# Potential use of Sodium Butyrate (SB) as an anti-virulence agent against *Vibrio cholerae* targeting ToxT virulence protein

**DOI:** 10.1101/2023.10.05.561138

**Authors:** Sushmita Kundu, Suman Das, Priyanka Maitra, Prolay Halder, Hemanta Koley, Asish K. Mukhopadhyay, Shin-ichi Miyoshi, Shanta Dutta, Nabendu Sekhar Chatterjee, Sushmita Bhattacharya

**Author notes:** **Corresponding author** Sushmita Bhattacharya Division of Biochemistry, ICMR- National Institute of Cholera and Enteric Diseases P-33, C.I.T. Road, Scheme XM, Beliaghata, Kolkata - 700 010, India.

## Abstract

Cholera, a diarrhoeal disease caused by gram-negative bacterium *Vibrio cholerae* remains a global health threat in developing countries owing to its high transmissibility and increase in antibiotic resistance. The current issue is to overcome the problem of resistance by antimicrobial therapy. There is a need for alternative strategies with an emphasis on anti-virulent approaches to alter the outcome of bacterial infections. *Vibrio cholerae* causes cholera by secreting virulence factors in the intestinal epithelial cells. Virulence factors help in cholera toxin production and colonisation during infection. Here, we show that sodium butyrate (SB), a small molecule, had no effect on bacterial viability but was effective in suppressing the virulence attributes of *V. cholerae*. The production of cholera toxin (CT) was downregulated in a standard *V. cholerae* El Tor strain and two clinical isolates when grown in presence of sodium butyrate. Analysis of mRNA and protein levels further demonstrated that sodium butyrate reduced the expression of the ToxT-dependent virulence genes like *tcpA* and *ctxAB*. DNA-protein interaction assays conducted at cellular (ChIP) and in *in vitro* conditions (EMSA) indicated that sodium butyrate weakens the binding between ToxT and its downstream promoter DNA, likely by blocking DNA binding. Furthermore, the efficacy of sodium butyrate was confirmed by showing its anti-virulence activity and tissue damage recovery in animal models. Collectively, these findings suggest that sodium butyrate (SB) has the potential to be developed as an anti-virulence agent against *V. cholerae* in place of conventional antibiotics or as an adjunctive therapy to combat cholera.

**IMPORTANCE:** The world has been facing an upsurge in cholera cases since 2021 with a similar trend continuing into 2022 with over 29 countries reporting cholera outbreaks (World Health Organization 16 December 2022 Disease Outbreak News; Cholera – Global situation). Treatment of cholera involves oral rehydration therapy coupled with antibiotics to reduce the duration of the illness. However, over the last few years, there has been indiscriminate use of antibiotics that contributed largely to the reservoir of antibiotic-resistant strains. In this study, we have addressed the problem of antibiotic resistance by targeting virulence factors. The screening of several compounds led to the identification of a small molecule, sodium butyrate that inhibits the virulence cascade in *V. cholerae*. We demonstrated that (i) sodium butyrate intervened with ToxT protein-DNA binding and subsequently affected the expression of ToxT-regulated virulence genes (*ctxAB* and *tcpA*) (ii) Sodium Butyrate is a potential therapeutic candidate for development of novel antimicrobial agents.

## INTRODUCTION

Cholera, an acute secretory diarrhoea is primarily caused by the Gram-negative, curved bacteria *Vibrio cholerae* through the ingestion of contaminated food and water. (1). The disease is marked by vomiting, copious amounts of non-bloody watery diarrhoea, severe loss of body fluids and electrolyte imbalance. The world is facing a drastic upsurge of the ongoing 7^th^ cholera pandemic since mid-2021 which is characterized by the occurrence of frequent outbreaks along with the increased incidence in mortality rate (2). This trend continued into 2022 and as of February 2023, at least 18 countries continued to report cases on cholera. The mortality linked to those outbreaks were particularly concerning as many countries were reporting higher CFR (case fatality ratios) than previous years (2). Despite the identification of approximately 200 *V. cholerae* serogroups, only the O1 (classical and El Tor biotypes) and O139 serogroups of *V. cholerae* have been linked to epidemic and pandemic cholera (3, 4).

The clinical symptoms of this disease is primarily induced by the production of two essential virulence factors, toxin-coregulated pilus (TcpA) and cholera toxin (CT) (5). TcpA, a major colonization factor in *V. cholerae* is responsible for microcolony formation and mediating bacterial attachment to intestinal epithelial cells (6). After microcolony formation, the bacteria start expressing cholera toxin (CT), an A-B_5_ exotoxin that is internalized by the host epithelial cells and accounts for severe diarrheal symptoms in this disease (7, 8). Co-ordinate expression of these virulence genes is directly under the control of the master regulator, ToxT, and given its pivotal role, numerous studies have been focused on its regulation, leading to the characterization of multiple mechanisms that contribute to its stringent regulation. ToxT belongs to the AraC/XylS family of transcriptional activators, which are characterized by the presence of two domains: an N-terminal domain that has been linked to effector binding and potential ToxT monomer association, and a DNA-binding domain at the C-terminus that has a homology to AraC/XylS (9). Activation of *toxT* in turn is carried out by protein complex present in the inner membrane, ToxRS and TcpPH (10). TcpPH is further activated by two activators AphA and AphB, which reacts to cell density, anaerobiosis, and other environmental factors (11, 12).

The mainstay therapy of cholera is an oral rehydration solution (ORS), which contains a variety of salts and glucose, to avoid dehydration (13, 14). Without intervention, the chance of surviving cholera can be as low as 50%; however, ORS supplementation reverses that chance to more than 99% (15). In some cases, antibiotics (16)are used as a secondary treatment as they can reduce the duration of the illness, however patients still run the risk of suffering from extreme dehydration caused by cholera toxin. Therefore, the use of antibiotics may not be a sustainable therapeutic option for cholera. Moreover, the intricacy of this issue is further increased by the emergence of antibiotic-resistant *V. cholerae* strains by the frequent acquisition of extra chromosomal mobile genetic elements from closely/distantly related bacterial species (17, 18). There are also vaccines against *Vibrio cholerae* but efficacy of vaccines is not 100% (19).The prevailing situation triggers the need for development of novel strategies that can specifically target the virulence factors of the pathogenic bacteria, leaving the large number of beneficial bacteria in the microbiome intact, thus limiting the unpleasant side-effects of current antibiotics.

Anti-virulence drugs are gaining popularity as they are combating diseases using different strategies unlike antibiotics. Firstly, they disarm the pathogen by targeting their virulence factors and then enabling the immune system to eradicate the infection (20). Secondly, this strategy circumvents several shortcomings of antibiotics as it enforces less pressure on emergence of resistant strains and reduce the risk of affecting commensal microbiota. Previous studies identified small molecules such as virstatin, toxtazin B, unsaturated fatty acids that target virulence gene regulatory cascade (15, 21, 22). Along with small molecules, several herbal products and bioactive compounds are also reported to be potent repressor of the virulence factors such as 6-Gingerol inhibiting the binding of CT to GM1(23), zinc oxide nanoparticles disrupting the secondary structure of CT (24), carbohydrate inhibitors (25) and fucosylated molecules (26) interfering with the activity of cholera toxin.

Short chain fatty acids such as butyrate are microbial metabolites synthesized from fermentation of dietary fibres in the colonic lumen. Multiple studies have reported the substantial effects of butyrate on host immunity, and it also hinders the colonizing ability of various pathogenic enteric bacteria in intestinal epithelial cells (27, 28). In colorectal cells, butyrate treatment is reported to induce a antimicrobial peptide, cathelicidin (29). In addition, the anti-microbial and anti-virulence activity of butyrate has been well characterized in a variety of pathogenic bacteria such as *Salmonella typhimurium, Clostridium perfringens* (30), *Escherichia coli* (31), *Staphylococcus pseudointermedius, Acinetobacter baumannii* (32), *Vibrio campellii* (33), and *Vibrio parahaemolyticus* (34).Despite extensive research on butyrate in recent years, there is still no report regarding the applicability of butyrate against *Vibrio cholerae*.

In this study, we have investigated the effect of sodium butyrate (SB) on the virulence factors of *Vibrio cholerae* with a particular emphasis on cholera toxin and adhesion factor TcpA production. This compound significantly reduced the expression of cholera toxin and TcpA in *in vitro* as well as *in vivo* conditions. Moreover, the potential molecular mechanisms behind the Sodium Butyrate mediated virulence gene inhibition in *V. cholerae* were also examined.

## RESULTS

### Viability of *V. cholerae* in presence of Sodium butyrate (SB)

The effect of sodium butyrate on the viability was examined on a standard *V. cholerae* (El Tor strain) and two clinical isolates (El Tor variant). The MICs of SB against strains N16961, Micro 78, BCH13298 were found to be 80mM and MBCs were found to be 160mM respectively. All the experiments were performed at sub-MIC concentrations of SB. In some experiments, MIC of SB was used to compare the effect with different treatment groups. The effect of SB on *V. cholerae* growth was investigated by counting the live cells on the plates in a time-dependent manner (Fig. 1). The growth rate of N16961 (Fig. 1A), Micro 78 (Fig. 1B) and BCH13298 (Fig. 1C) were measured in LB medium along with exogenously added different doses of SB. The result indicated that there was a complete inhibition of *V. cholerae* growth at 2xMIC (160 mM). No bacteria were recovered after 12 h in N16961 at 2xMIC. The bactericidal activity of SB was much more potent in strains BCH13298 and Micro78 as no bacteria were recovered after 6 h in BCH13298 and after 8 h in Micro78 at 2xMIC. At 1xMIC (80mM), the growth pattern of all the three strains were almost similar with the viable count reduced to 65% 2 h post-initial inoculum. Within the initial 8 h, the viability of the bacteria was further reduced to 20% and from 8 to 30 h, there was a bacteriostatic effect on the remaining viable bacteria. At sub-MICs (10, 20 and 40 mM) there was no significant difference in growth rate as compared to the untreated control. These results indicate that sub-MIC concentrations of sodium butyrate has no effect on viability of *V. cholerae* strains.

**Fig. 1).**
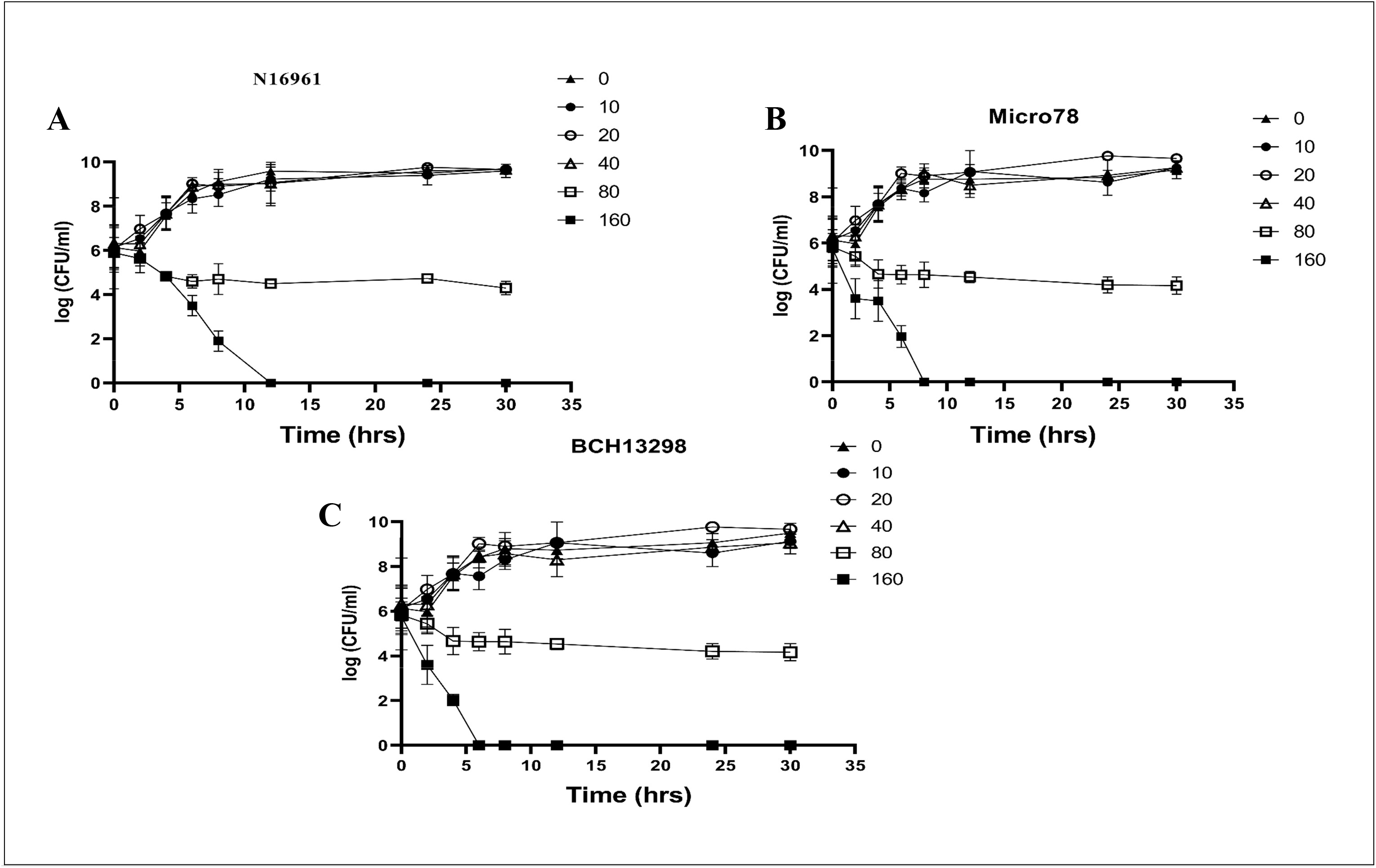
Effect of Sodium butyrate (SB) on the growth of *Vibrio cholerae*. Effect of SB on *Vibrio cholerae* growth at MIC and sub-MICs. El Tor *V. cholerae* strains were grown in LB broth along with SB at 2xMIC (160 mM), SB 160 (▪); 1xMIC (80 mM), SB 80 (□); 1/2^th^ MIC (40 mM), SB 40 (Δ), 1/4^th^ MIC(20 mM), SB 20 (**○**),1/8^th^ MIC(10mM),SB 10 (●) along with the untreated control, SB 0 (▲) for the indicated time periods (2, 4, 6, 8, 12, 24 and 30 h). The viable bacterial counts in CFU per ml detected by the plate count method were represented graphically. **(A)** wild-type *V. cholerae* N16961, **(B)** multidrug resistant strain Micro78 and **(C)** multidrug resistant strain BCH 13298. All data are represented as mean ± S.E.M. (*n* = 3).

### Inhibition of Cholera toxin production

Host pathogen interaction is dependent on cholera toxin production and TCP expression. To begin with, El Tor *V. cholerae* strain N16961 and two multidrug resistant El tor variant strains Micro 78 and BCH13298 were used for the study to see the effect of sodium butyrate (SB) on cholera toxin production (CT) (Fig. 2). We explored the classical GM1-CT ELISA assay to detect the level of CT secreted in the supernatant of the bacterial culture grown in presence or absence of SB. The sub-MIC concentrations of SB that had no adverse effect on viability of the bacteria was found to have significant effect on the CT production. There was almost 4-fold, 3-fold and 6-fold reduction in cholera toxin (CT) production at 20 mM (1/4^th^MIC) for N16961(Fig. 2A), Micro 78(Fig. 2B) and BCH13298(Fig. 2C) respectively. Similarly, at 40 mM (1/2^th^MIC) SB there was 12-fold, 7-fold and 10-fold reduction in CT in N16961(Fig. 2A), Micro 78(Fig. 2B) and BCH13298(Fig. 2C) as compared to the SB-free culture. The results obtained from this assay showed that although sodium butyrate(SB) had no effect on bacterial growth, but it demonstrated significant activity in reduction of CT production.

**Fig. 2).**
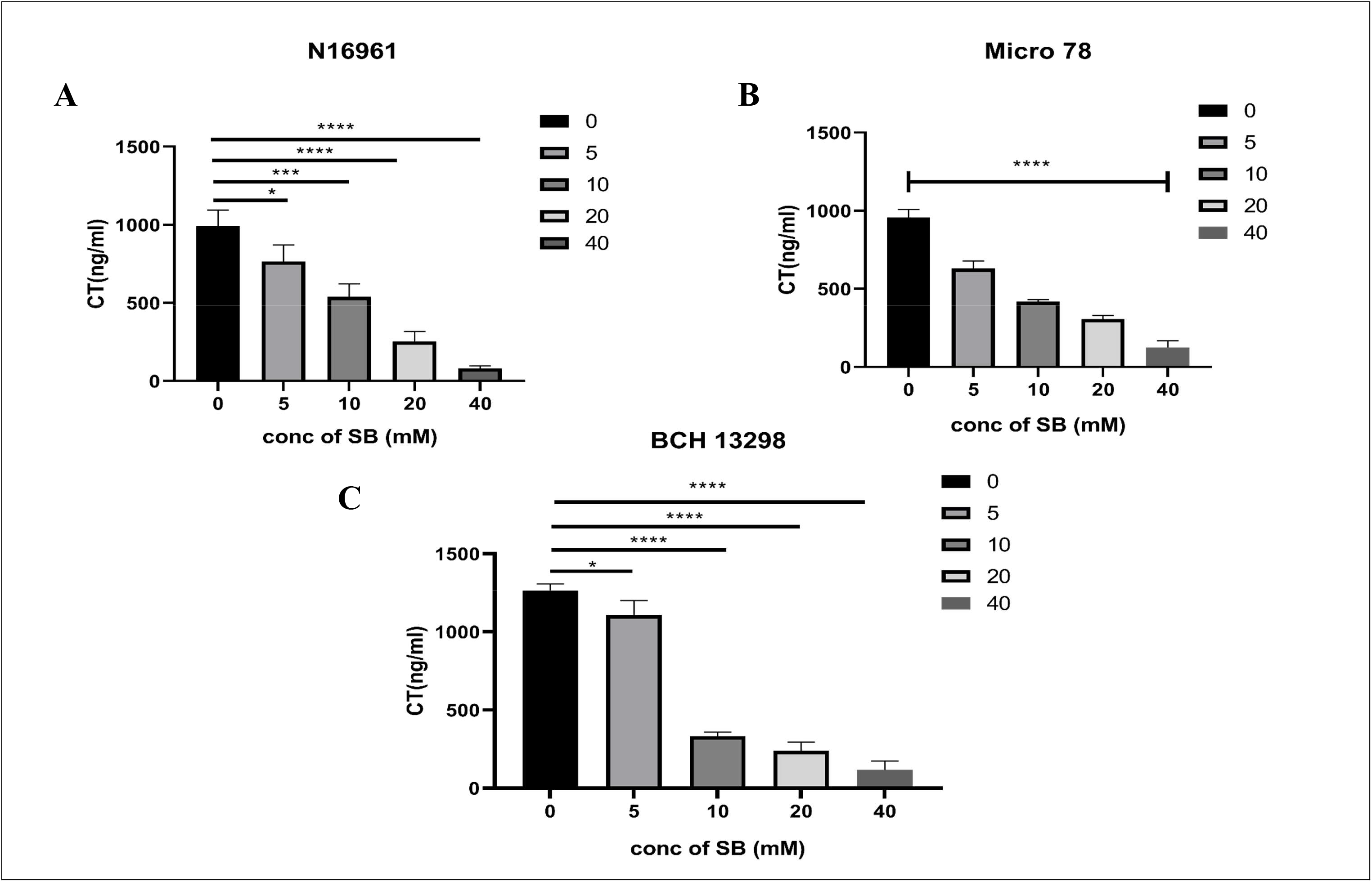
Dose-dependent inhibitory effects of Sodium butyrate (SB) on cholera toxin (CT) production in different *Vibrio cholerae* O1 El Tor strains. *In vitro* CT production from different O1 El Tor strains were measured by ELISA from the samples grown in AKI media (pH=7.5, 37°C, static condition for 18 hours) with (5(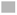), 10(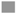), 20(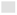) and 40(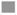) mM SB) or without SB respectively. The amount of CT produced is graphically represented **(A)** *V. cholerae* N16961 **(B)** Micro78 and **(C)** BCH 13298. All data are represented as mean ± S.E.M. (*n* = 3). One-way ANOVA was performed. Significance levels were denoted as ∗for *P <* 0.05, ∗∗∗ for *P <* 0.001, ∗∗∗∗for *P <* 0.0001.

### Effect of sodium butyrate (SB) on virulence-related gene expression

To understand the underlying mechanism behind CT reduction, we further checked the transcript levels of virulence genes. The inhibitory effect of SB on CT production was analysed using El Tor strain N16961 by assessing *ctxA*, *ctxB* gene transcripts and other virulence genes by qRT-PCR (Fig. 3A). There was almost 4.3-fold and 3.8-fold reduction in *ctxA* and in *ctxB* transcripts respectively. In addition to this, the influence of SB on transcription of *toxS, toxR, tcpH, tcpP, tcpA, toxT* genes were also analysed. Along with *ctxAB* transcripts there was an impressive reduction of 9-fold in *tcpA* transcripts. However, there was no effect of SB on the transcription of *toxS, toxR, tcpH, tcpP* and *toxT*. The results obtained from this assay indicated a significant (p<0.0001) inhibitory effect of SB on the transcription of *ctxAB* and *tcpA* genes, however, no difference in the transcript level of *toxT* gene was observed in the presence of SB.

**Fig. 3).**
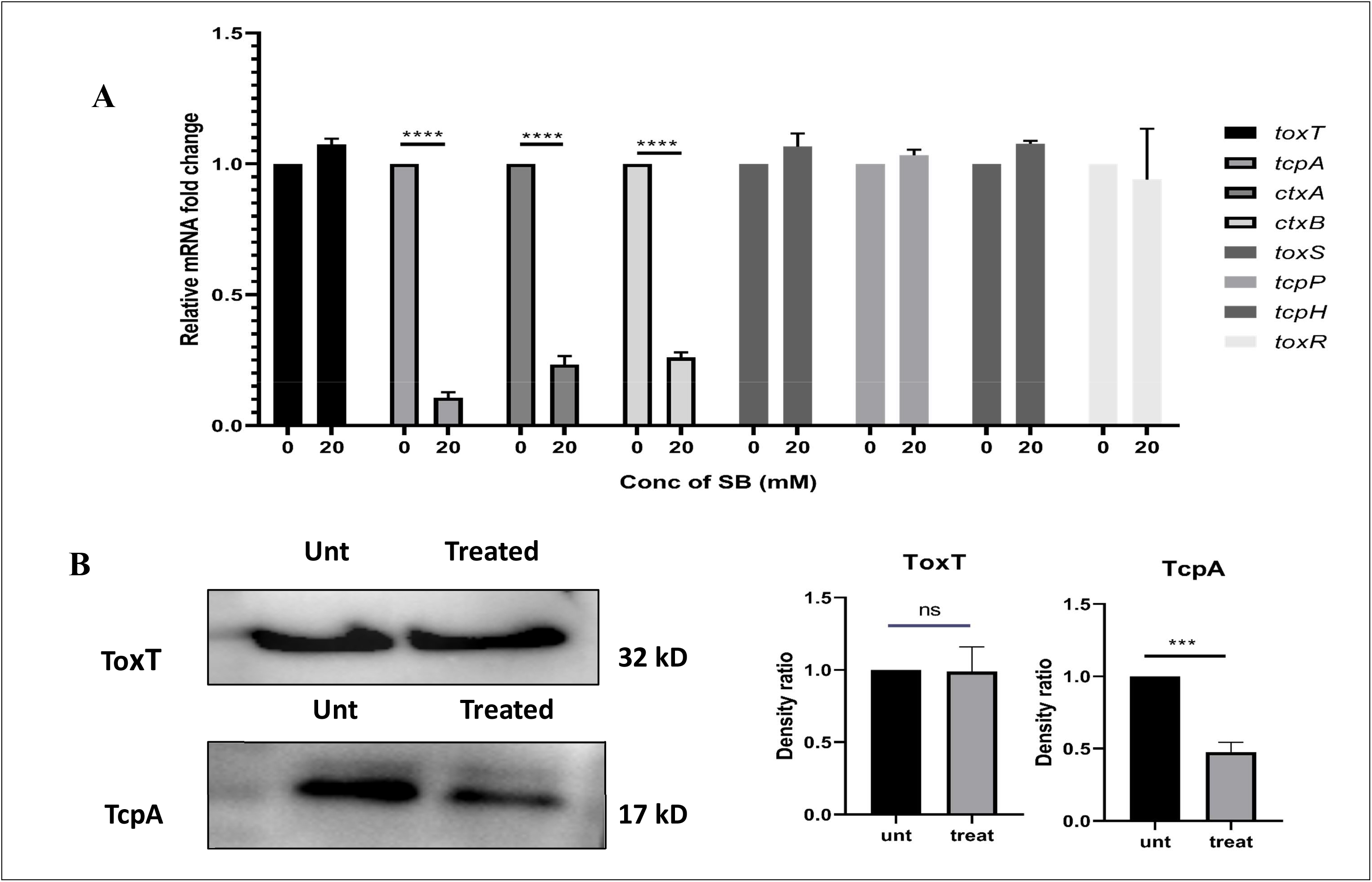
The effect of Sodium butyrate (SB) on virulence attributes of *Vibrio cholerae* O1 El Tor strain N16961. *Vibrio cholerae* N16961 was cultured under virulence inducing conditions (AKI media [pH 7.5], 37°C, static condition for 18 hours) in the presence or absence of 20mM SB. **(A)** Transcript levels of *ctxA*(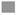), *ctxB*(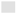), *toxS*(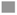)*, toxR*(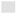)*, tcpH*(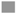)*, tcpP*(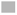)*, tcpA*(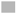)*, toxT*(▄) virulence genes were analyzed by real-time PCR and normalized using *recA* gene as the internal control and graphically represented. One-way ANOVA was performed. (**B)** Expression of major virulence proteins ToxT and TcpA were determined by western blotting followed by densitometric analysis which is graphically represented. Students t-test was performed. All data are represented as mean ± S.E.M. (*n* = 3). Significance levels were denoted as n.s for non-significant, ∗∗∗ for *P <* 0.001 and ∗∗∗∗ for *P <* 0.0001.

To rule out the possible mechanism that the reduced transcription of *ctxAB* and *tcpA* genes depend on the post-transcriptional or translational regulation of *toxT*, the levels of ToxT and TcpA proteins were measured. To this end, western blot assay was performed on total protein extracts obtained from the treated and untreated samples (Fig. 3B). The results demonstrated that SB had no relevant effect on the cellular levels of ToxT. Moreover, the reduction in TcpA protein level in presence of SB was consistent with the corresponding transcriptional profile. These observations are noteworthy in indicating that inhibition of SB is related to virulence cascade but not with ToxT production.

### Sodium Butyrate inhibits the binding activity of ToxT to its downstream promoter at cellular level

The results obtained from the qRT-PCR and western blot data showed a stable production of ToxT protein in presence or absence of the compound. Since ToxT is the direct transcriptional activator of *tcpA* and *ctxAB*, there can be a possibility that this compound affects the transcriptional activity of ToxT thereby reducing the activation of *tcpA* and *ctxAB* genes. To clarify the interaction between ToxT protein to promoter region of *tcpA* gene in presence of SB under physiological conditions where ToxT co-exist with other transcriptional factors, we performed ChIP assay. As shown in (Fig. 4A and B), ToxT occupancy at *tcpA* promoter region was drastically reduced in presence of SB suggesting a strong interference on the binding activity of ToxT to its specific DNA site at the cellular level. Hence, SB acts on the transcriptional activation of virulence factors by inhibiting ToxT-DNA binding.

**Fig. 4).**
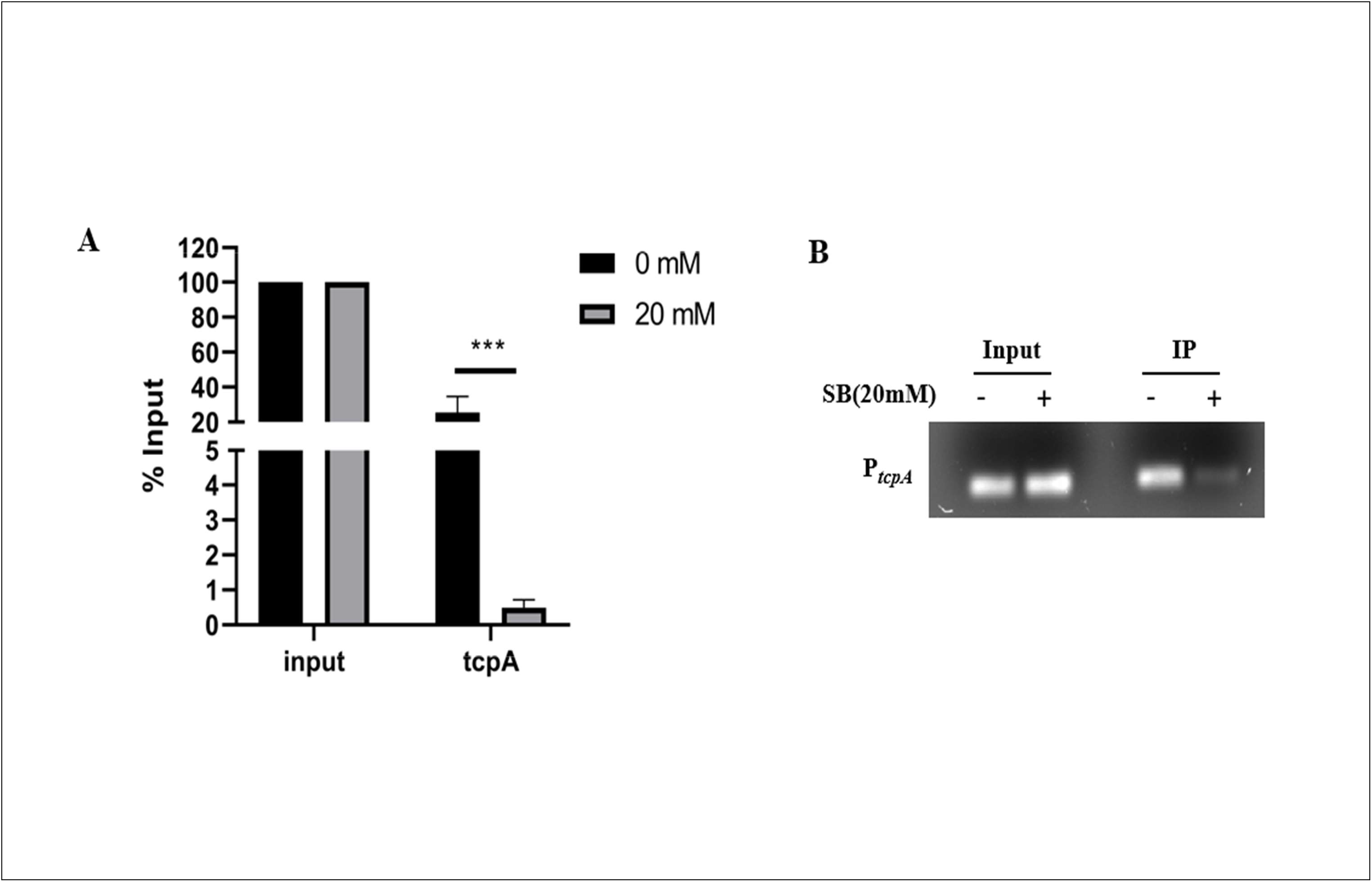
Sodium butyrate (SB) prevents the ToxT-promoter binding to virulence genes. *V. cholerae* cells were grown in presence or absence of SB for 4h. The culture was crosslinked in 1% formaldehyde and the assay was performed with or without anti-ToxT antibody (input). **(A)** Expression of *tcpA* was checked and compared to input in real-time PCR assay for 0mM SB (^▄^) and 20mM SB (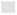) treated cells. The data were represented graphically and compared to input; two-way ANOVA was performed significance. IP, Immunoprecipitation; Levels were denoted as ∗∗∗∗ for *P <* 0.0001. **(B)** Promoter DNA IP with anti-ToxT antibody was determined by agarose gel electrophoresis.

### Sodium Butyrate (SB) reduces ToxT-DNA binding affinity *in vitro*

Based on the results obtained from the ChIP assay, it led to the hypothesis that SB is interfering the binding of ToxT to the DNA. To further confirm this hypothesis, we expressed, purified ToxT proteins and performed *in vitro* electrophoretic mobility shift assay (EMSA). ToxT protein was expressed in pHIS-Tev plasmid and furthermore the His-tag protein was purified by Ni-NTA purification (Fig. S1). EMSA revealed the increase in DNA-ToxT complex formation in a ToxT dependent manner. When 20 mM of SB was added, reduction in DNA-ToxT complex formation was observed (Fig. 5A). Moreover, on adding increasing amounts of SB there was a concentration-dependent suppression of ToxT binding to the DNA, and the formation of the DNA-ToxT complex was completely abolished at 40 mM (Fig. 5B). In contrast, when another butyrate derivative Tributyrate (TB) was used, it showed no inhibition on ToxT-DNA binding (Fig. 5C).

**Fig. 5).**
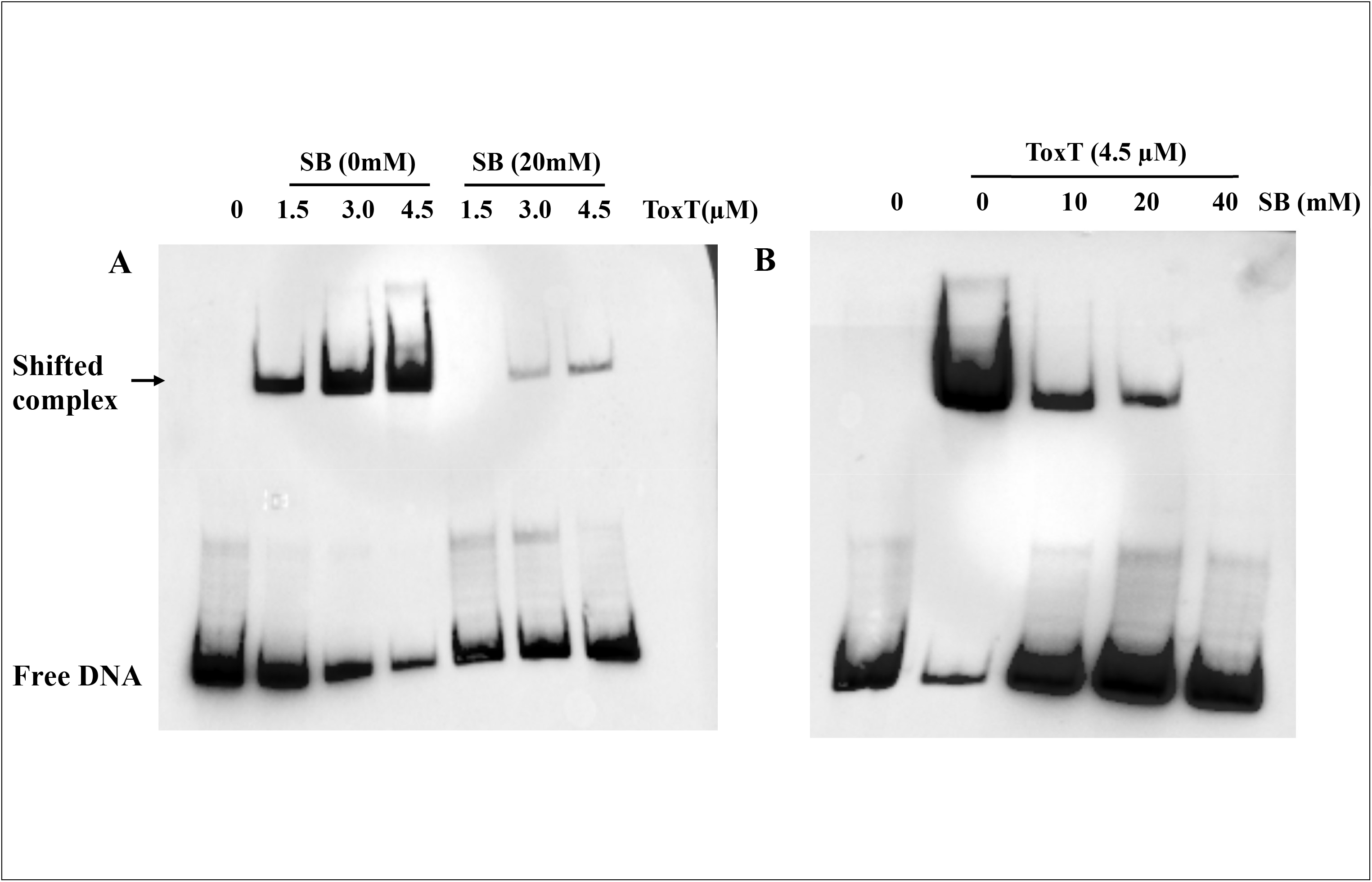
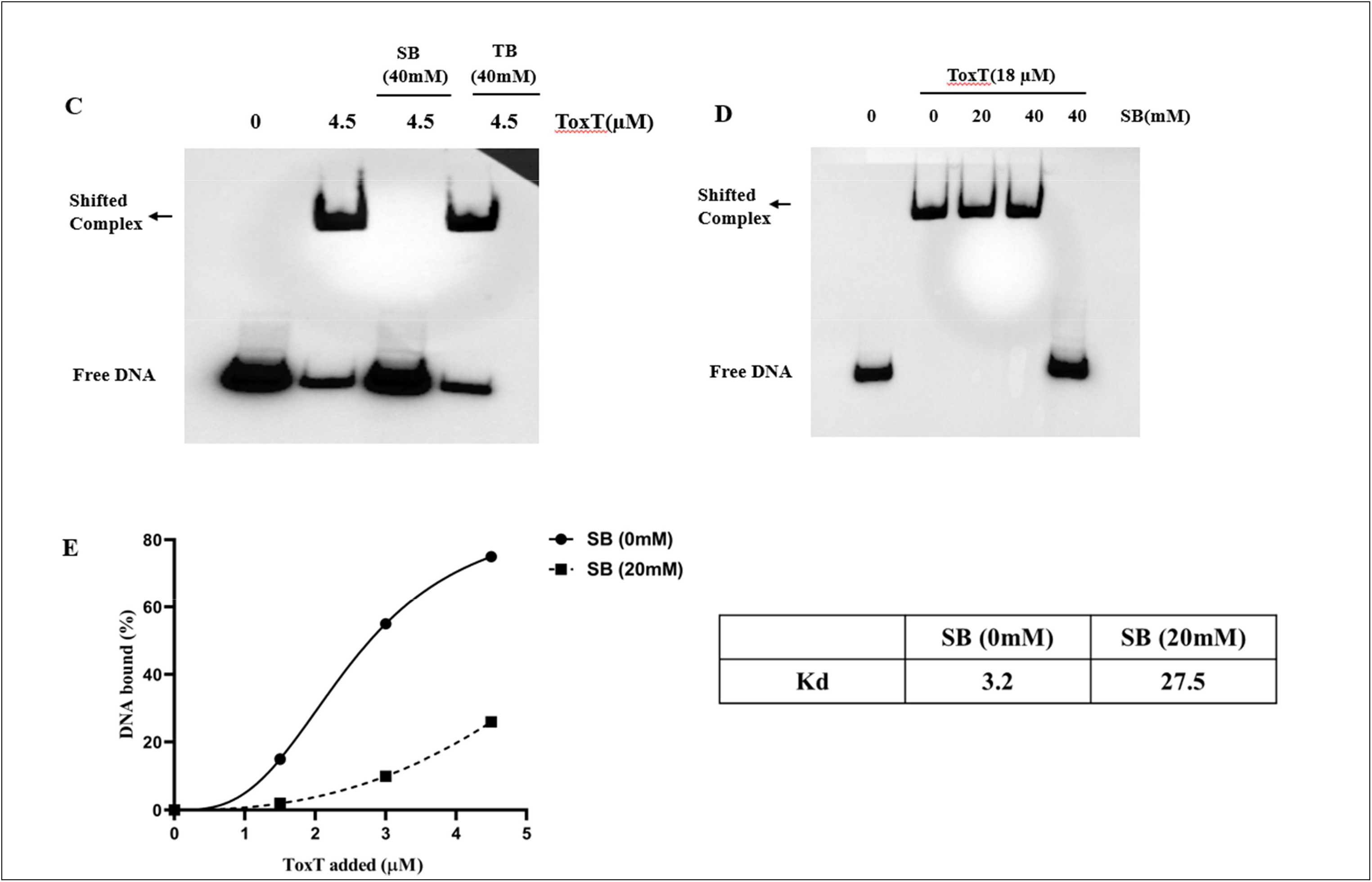
The effect of Sodium Butyrate (SB) on promoter binding activity of ToxT to *P_tcpA_* by EMSA. Gel mobility shift assay was carried out with the biotinylated labelled *tcpA*. promoter fragment (-123 to +28) combined with purified (His)6-ToxT protein. (**A)** Increasing concentration of ToxT protein (0-4.5 µM) was incubated with 20mM SB or 0mM of SB followed by the addition of P***_tcpA_*** DNA (**B)** The highest concentration of ToxT protein (4.5 µM) was incubated with increasing concentrations of SB (0-40mM) followed by the addition of P***_tcpA_*** DNA. (**C)** The highest concentration of ToxT protein (4.5 µM) was incubated with 40mM Sodium Butryrate (SB) or 40mM Tributyrate (TB) followed by the addition of P***_tcpA_*** DNA. The complexes were separated by electrophoresis on a 4% non-denaturing polyacrylamide gel. **(D)** Biotin labelled P***_tcpA_*** DNA (0.5µM) was incubated with SB (20 and 40mM) followed by ToxT protein (18 µM) addition and proceeded as described above. **(E)**The relative affinities of ToxT protein in presence or absence of SB was compared using the data from panel (A). The percentage of bound DNA was calculated and plotted against the concentration of ToxT added. The corresponding dissociation constant (K_d_) values were obtained from the graph.

To investigate the alternative hypothesis that SB can interact with DNA and inhibit the DNA-protein binding, P***_tcpA_*** DNA was incubated with SB followed by the addition of purified ToxT protein. No inhibition on DNA binding activity was observed under this reaction condition. These results reduced the possibility of our alternative hypothesis that SB prevents the ToxT-DNA interaction by binding with DNA (Fig. 5D). The K_d_ for ToxT in absence of SB was found to be 3.2 µM, while in presence of 20mM of SB it was increased to 27.5 µM (Fig. 5E), indicating a significant effect on the equilibrium between DNA-bound ToxT and free ToxT proteins. Together, these results confirm that SB specifically binds to ToxT and inhibits its binding to the promoter of *tcpA*.

### Molecular docking shows stable binding of Sodium Butyrate (SB) with ToxT

In addition to our *in vitro* results, we performed in silico binding to show the possible interacting residues of ToxT to sodium butyrate. The activator protein ToxT was molecularly docked with sodium butyrate to explore if any interaction exists between them. The 3D image of the receptor (ToxT) and ligand (sodium butyrate) after optimal docking as visualized is represented (Fig. 6A and B). The results revealed a stable interaction between ToxT protein and SB with a binding energy of -7.68 Kcal mol^-1^. From Table 6C, the key amino acids of ToxT protein that interacted with SB were Gln8, Thr85, Ser227, Ser223, Lys31 and Lys230. Previous studies have shown that the carboxylate head of UFAs (unsaturated fatty acids) were found to have interaction with the lysine residues Lys31, Lys230 and this interaction contributed to the allosteric regulation of ToxT activity (35). From our interaction study, we also obtained similar interaction between the lysine residues of ToxT protein and the carboxylate head of SB. This led to the hypothesis that SB might interact with the ToxT protein and form a stable complex which is further preventing the binding of ToxT to its target DNA site.

**Fig. 6).**
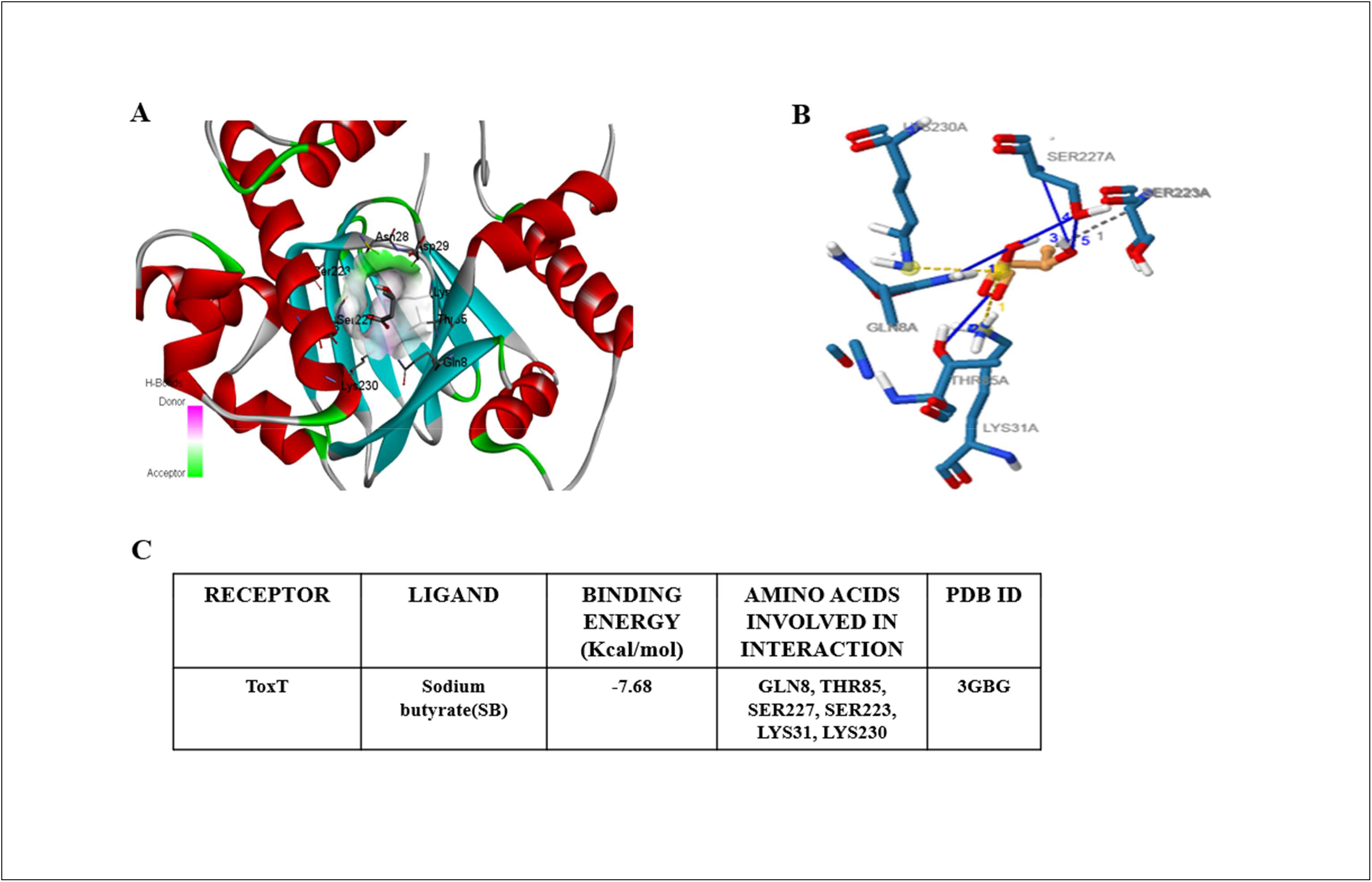
Binding of Sodium butyrate (SB) with ToxT *in silico*. Residues interacting with ligand(Sodium Butyrate) and receptor (ToxT) are represented in **(A)** 3D conformation and **(B)** 2D conformation. **(C)** Table showing the probable interacting residues and the binding energy.

### Adhesion to intestinal epithelial cells is affected by Sodium butyrate (SB)

Adhesion of *V. cholerae* to epithelial cells is facilitated by TcpA, an important colonization factor that is involved in the microcolony formation and adherence to the epithelial cells. The adhering ability of *V. cholerae* to the intestinal epithelial cells was assessed in HT-29 cell line as well as confirmed in *in-vivo* animal models.

Firstly, we observed the effect of SB on the adhering ability of *V. cholerae* in HT-29 cells (Fig. 7A and B) and further validated in rabbit ileal loop model (Fig. 7C and D) from the recovered viable bacteria. In the absence of SB, almost 10.5% and 15% bacteria adhered to HT-29 and rabbit intestinal cells respectively. However, in presence of higher doses of SB (40 and 80 mM) approximately 6% and 4% bacteria out of total viable bacteria were found to adhere to HT-29 cells and rabbit intestinal cells respectively. Therefore, the effect of SB on *V. cholerae* is more pronounced in *in-vivo* treatment compared to the *in-vitro* conditions. Additionally, the potential toxic effect of SB on HT-29 cells was also examined (Figure S2). SB treatment for 24 hours at varying concentrations (5-80mM) did not exhibit significant toxicity.

**Fig. 7).**
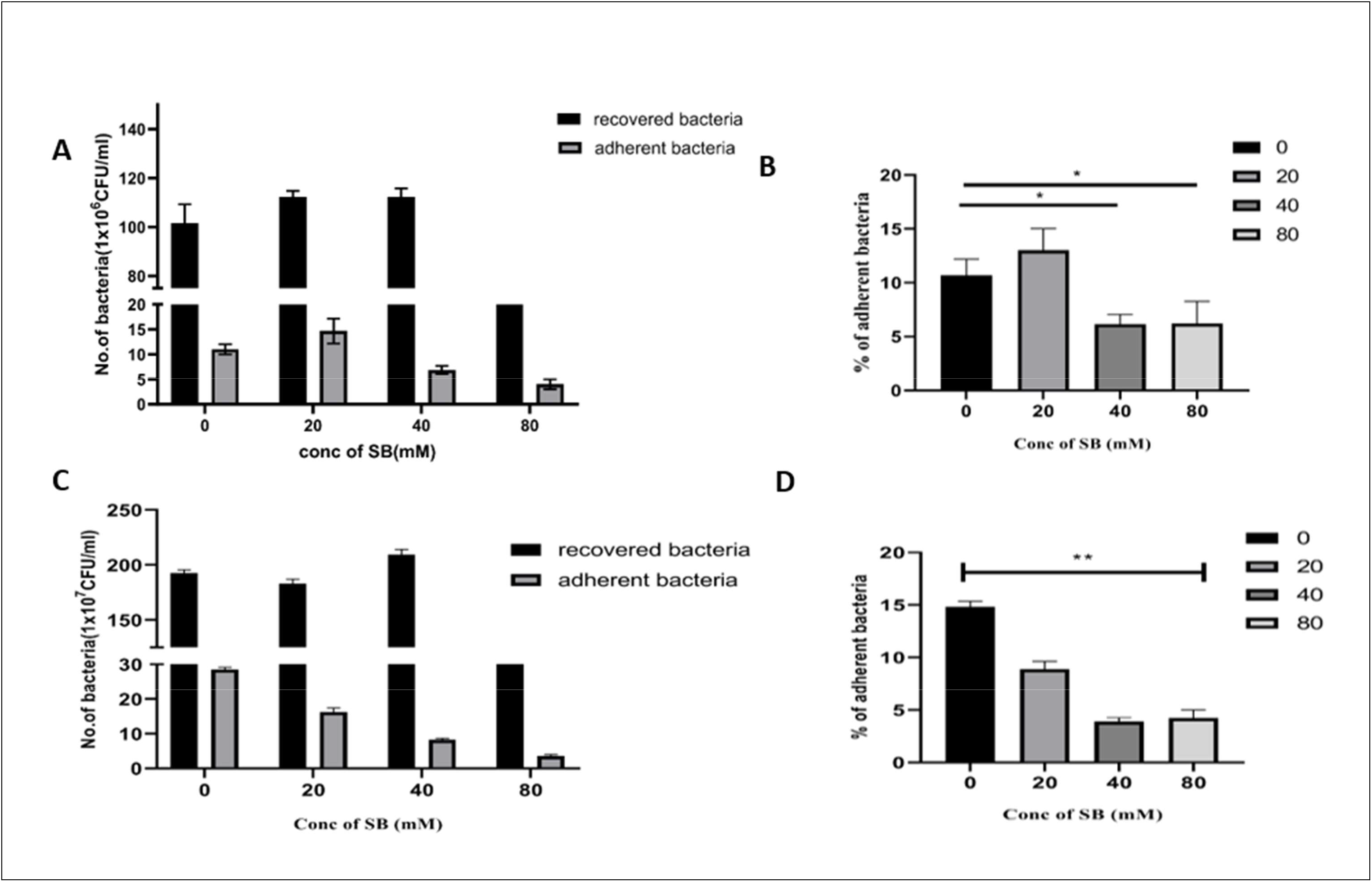
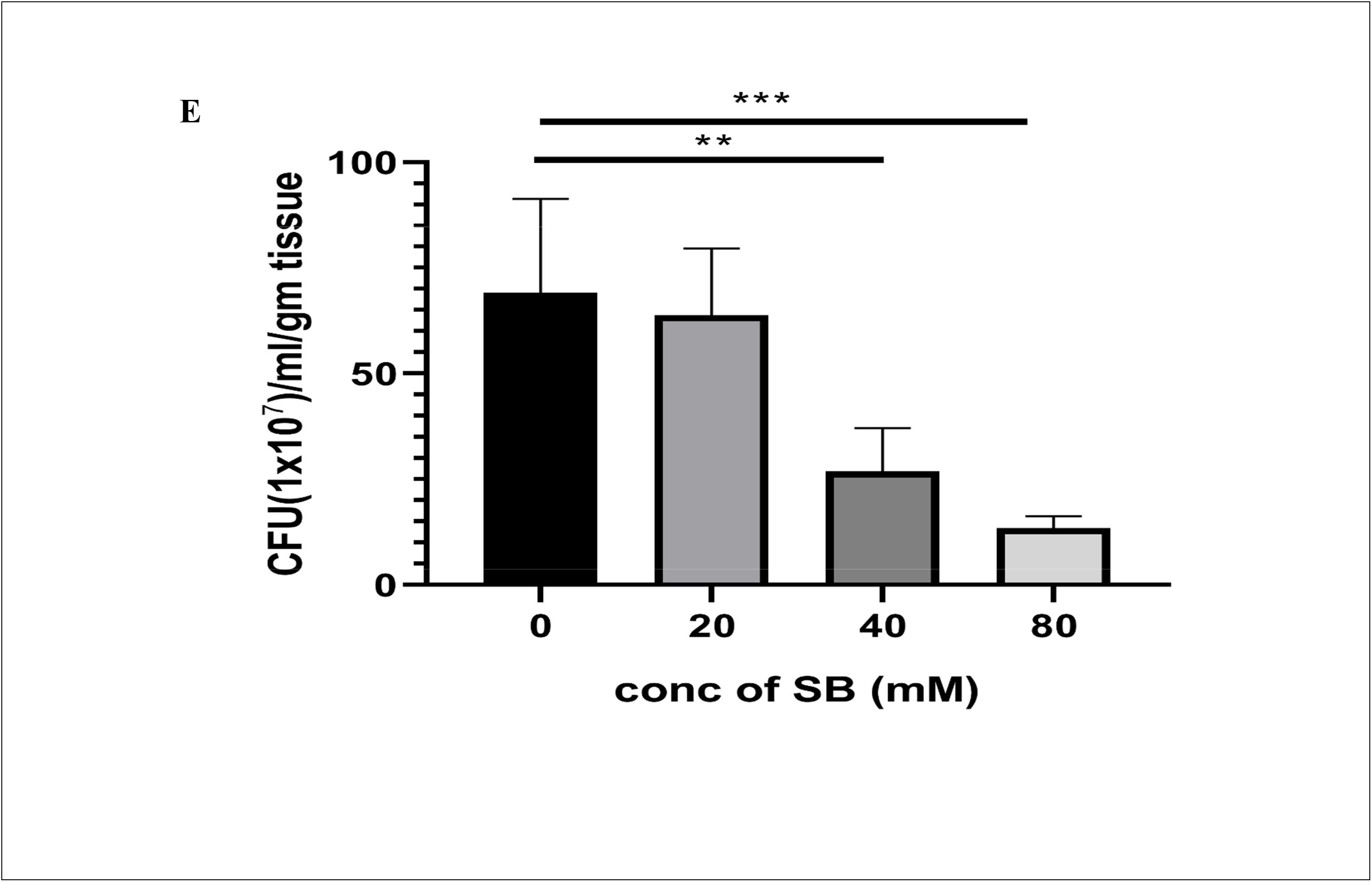
Influence of Sodium butyrate (SB) on the adhesion of *V. cholerae* to the epithelial cells. Different concentrations of SB at 0, 20, 40 and 80 mM were used to treat HT 29 cells. **(A)** Total number of recovered and adhered bacteria in each of the HT-29 culture samples are represented. **(B)** % of adhered *V. cholerae* to the HT-29 cell line. For rabbit ileal loop model, each loop was injected with 1×10^9^ CFU per ml *V. cholerae* N16961 with or without SB (at 20, 40 and 80mM) and loop injected only with PBS served as negative control loop. After 18 h, the animals were euthanized, and the loops were removed. **(C)** Total number of recovered and adhered bacteria to the rabbit ileal loop tissue samples is represented. (**D)** % of adhered *V. cholerae* to the rabbit ileal loop tissue samples. **(E)** Effect of SB on bacterial colonization in 4 to 5-day old suckling BALB/c mice. The mice were orogastrically challenged with 1x 10^5^ CFU/ml of *V*. *cholerae* strain N16961 in presence of different concentrations of SB or absence of SB and kept at 30°C for 18 hours without their mother. Bacterial colonization was calculated as CFU/ml/gm and graphically represented. One-way ANOVA was performed. Significance levels were denoted as ∗for *P <* 0.05, ∗∗for *P <* 0.01, ∗∗∗ for *P <* 0.001.

Next, we checked the colonization ability in suckling mice (Fig. 7E). A significant (p<0.001) reduction in colonization was observed at higher doses of SB (40 and 80mM). The colonizing ability of *V. cholerae* was decreased by more than 3-fold at higher doses of SB when compared to the untreated *V. cholerae* mice. These results suggest that SB inhibits the colonization ability of *V. cholerae* in suckling mice.

### *Vibrio cholerae* showed less CT production and fluid accumulation upon Sodium butyrate (SB) treatment

Next, we assessed the effect of SB on toxin production and fluid accumulation in rabbit ileal loop model as fluid accumulation is a symptom manifested during *Vibrio cholerae* infection. The effects of SB on fluid accumulation (Fig. 8A, B) and CT production (Fig. 8C) were assessed by using the rabbit ileal loop model. Varying concentrations of SB were administered separately into the ileal loop along with the bacterial inoculation (1×10^9^ CFU/ml). The effect of SB on fluid accumulation (F/A ratio) was evaluated. There was a marked reduction in fluid volume in loops that received the highest concentration of SB. The FA ratio of SB treatment loops (40 and 80 mM) showed more than 20-fold reduction in fluid accumulation as compared to the untreated loop (Fig. 8B). Experimental results suggested that the concentration of CT was significantly reduced at 18 hours post-administration for each of the SB concentrations tested. The CT level in the loops that received the highest concentration of SB (40 and 80mM) was reduced by more than 10-fold as compared to the CT level in the loop that received no SB (Fig.8C). Further, we have studied the real time PCR analysis of virulence genes, where we found that the expression of *ctxA* and *ctxB* gene was downregulated by more than 6-fold and 5-fold respectively at 40 and 80 mM respectively. In addition, another important virulence factor *tcpA* was also downregulated by 5-fold (Fig. 8D). These data demonstrate that SB effectively inhibited CT production in the rabbit small intestine.

**Fig. 8).**
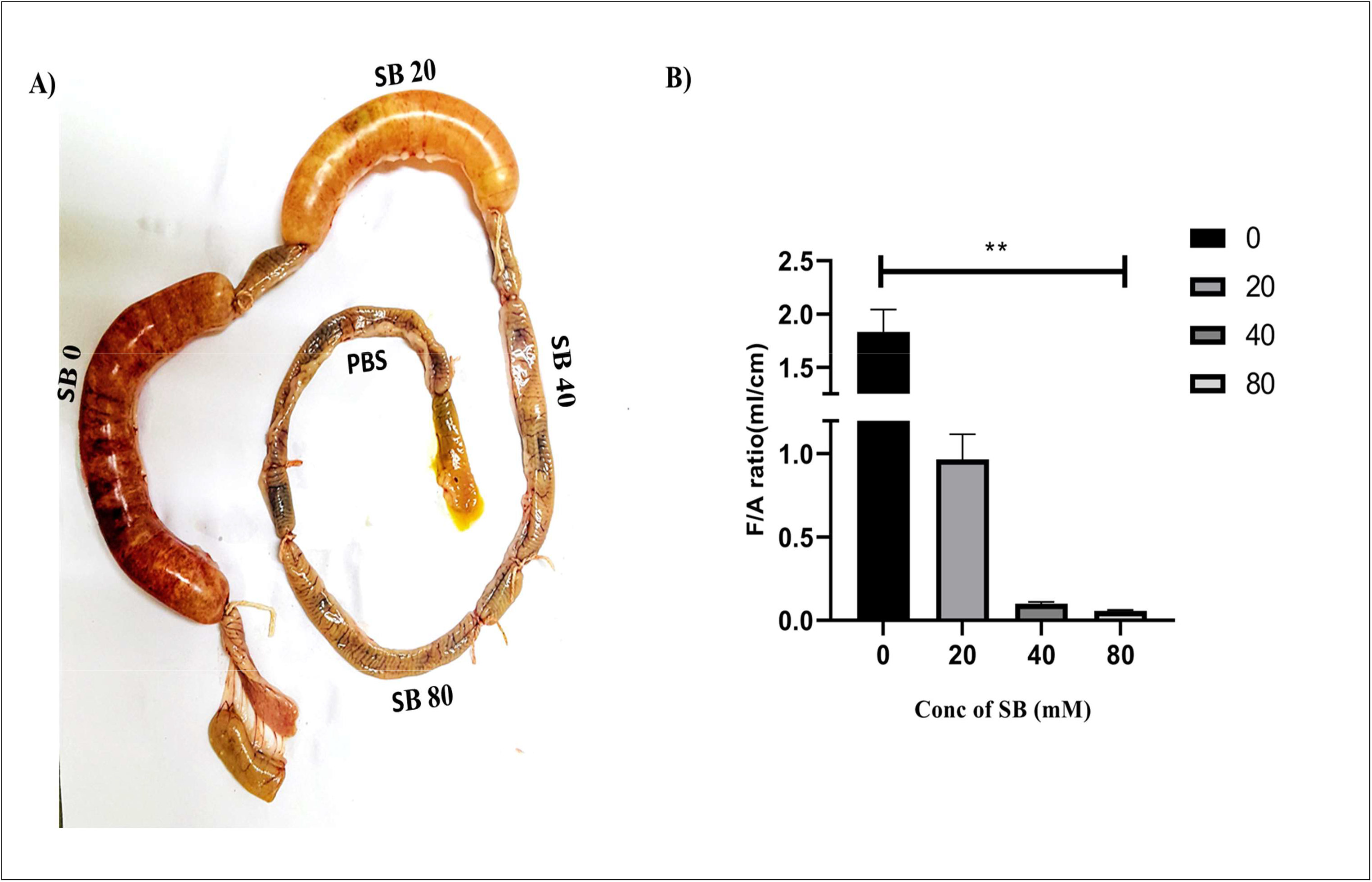
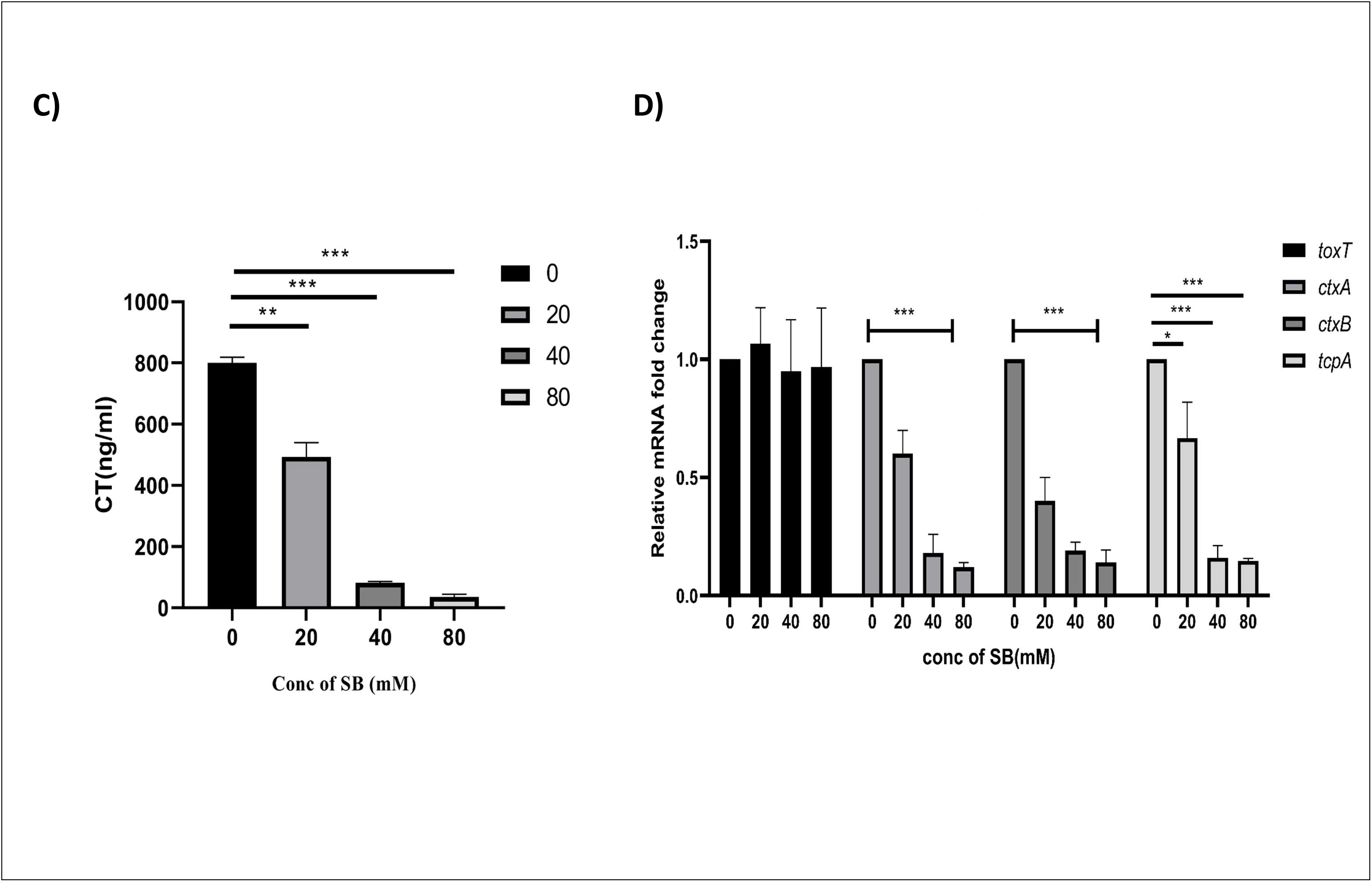
Inhibitory effect of Sodium butyrate (SB) on fluid accumulation, cholera toxin (CT) production and virulence gene expression. Each loop was injected with 1×10^9^ CFU per ml *V. cholerae* N16961 with or without SB (at 20, 40 and 80mM) and loop injected only with PBS served as negative control loop. After 18 h, the animals were euthanized, and the loops were removed. **(A)** Representative image of recovered rabbit intestine segment with infected loop at 18 h after infection. **(B)** Fluid accumulation per unit area (F/A) ratio of each loop was evaluated and graphically represented as mean ± S.E.M. (*n* = 3). **(C)** The accumulated fluid samples from SB treated and untreated ileal loops were used to measure the CT production by ELISA. (**D)** Relative expression of major virulence genes *ctxA* (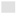), *ctxB* (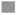), *tcpA* (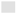), *toxT* (▄) were analyzed by real-time PCR. The normalization was done using *recA* gene as the internal control. One-way ANOVA was performed. All data are represented as mean ± S.E.M. (*n* = 3). Significance levels were denoted as ∗for *P <* 0.05, ∗∗for *P <* 0.01, ∗∗∗ for *P <* 0.001.

These results strongly imply that administration of SB *in vivo* could considerably lessen the amount of CT production and secretory diarrhoea, which should also reduce the severity and duration of disease in cholera patients.

### Treatment with SB rescues rabbit intestinal tissue damage caused by infection with wild type *V. cholerae* N16961

The results of the rabbit intestinal fluid-accumulation (RIL) assay prompted us to look into the microscopic alterations of the intestinal tissues. Development of therapeutics are dependent on tissue damage recovery in pre-clinical studies. Histopathological analysis of the rabbit ileum showed that all the layers of mucosa were severely damaged by the wild *V. cholerae* N16961 infection. There was a gross alteration in structure with deformed villi and necrotic changes in mucosa, sub-mucosa and lamina propria. Haemorrhage at the site of muscularis mucosa was also seen (Fig.9A). In contrast to that, SB treatment (20, 40 and 80mM) completely recovered the damage caused by infection as normal villi and mucosal structure was noticed (Fig.9 B,C,D). The ileal loop tissue treated with PBS displayed normal villi structure and did not exhibit any alteration to the intestinal mucosa (Fig.9E). Hence, it is confirmed that sodium butyrate inhibits diarrhoea and at the same time also helps in recovery of tissue damages.

**Fig. 9).**
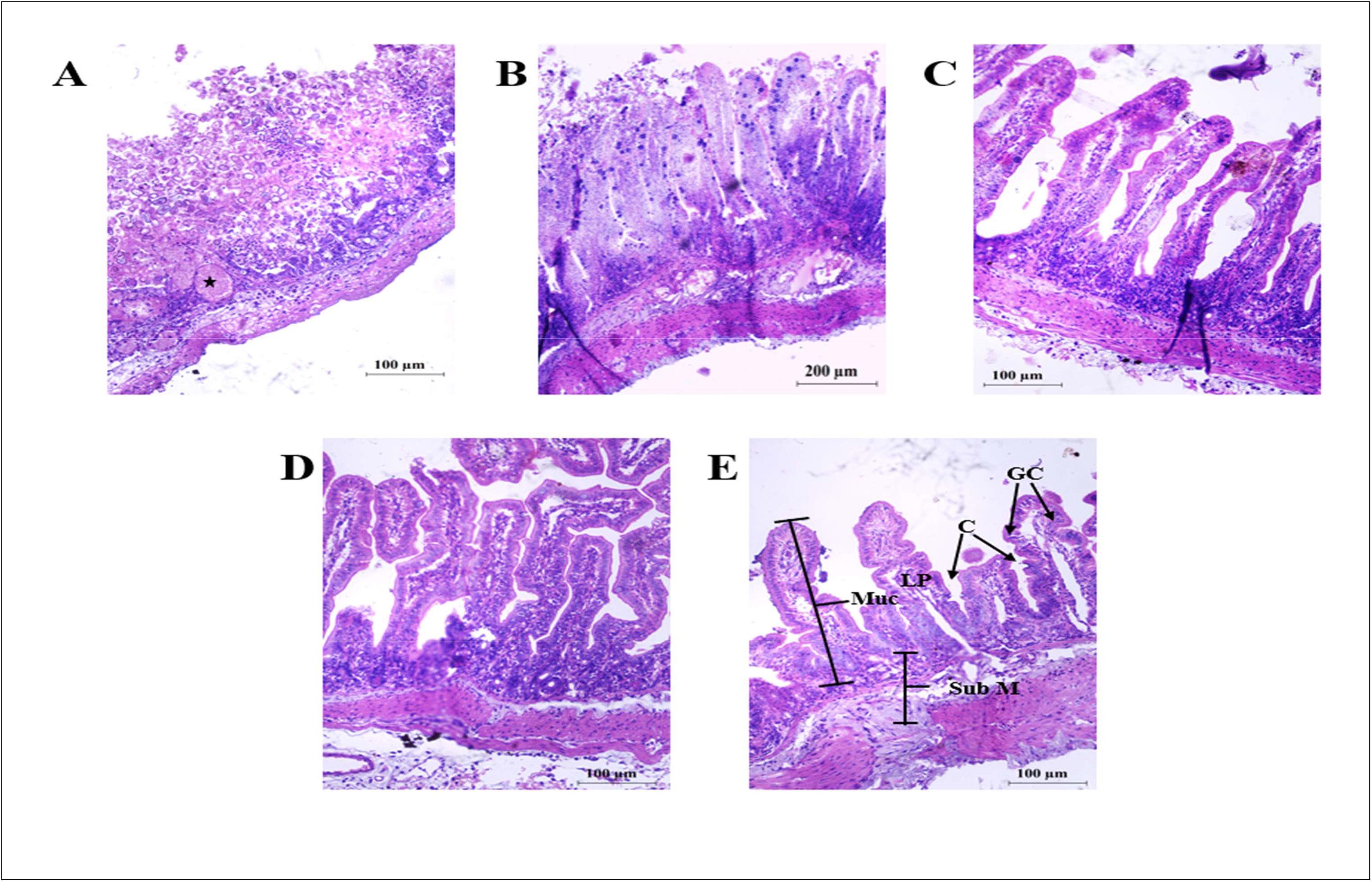
Histological staining of the rabbit intestine showed that Sodium Butyrate (SB) restores the damage caused by *V. cholerae* N16961 to the intestinal lining. Rabbit ileal loops challenged with (A) *V. cholerae* infection (1×10^9^ CFU per ml). (B) *V. cholerae* with 20 mM of SB. (C) *V. cholerae* with 40 mM of SB. (D) *V. cholerae* with 80mM SB. **(E)** PBS were subjected to histopathological study (H&E). Severe mucosal damage was observed including disruption to villus structure and necrotic changes in mucosa, sub-mucosa and lamina propria in infected ileal section which was restored upon SB treatment.; GC, Goblet Cells; C, Crypts LP, Lamina Propria; Muc, Mucosa; SubM, Submucosa, LP, Lamina Propria *, indicating the site of haemorrhage.

## DISCUSSION

The conventional methods of treating bacterial infections rely on the use of antibiotics that reduce bacterial viability. However, if viability is inhibited, it will inevitably give rise to antibiotic resistant strains. The rise of antibiotic-resistant bacteria along with the reduced effectiveness against the majority of present antibiotics has become a worldwide problem to public health and this is one of the major reasons for the growing healthcare costs. So, there is an imminent need for new treatment therapies that can impede the harmful bacteria without affecting the normal microflora of the gut (36, 37). An emerging goal to combat bacterial infection in this post-antibiotic era involves the use of molecules that can target important virulence factors rather than affecting the bacterial growth. After screening several compounds, Sodium butyrate (SB), a small molecule, was selected for this study as it was found to possess diverse biological activities on various pathogenic organisms.

In this present study, we examined the sub-inhibitory concentrations of SB on *V. cholerae* to access its wide range of biological functions under pathogenic conditions. According to our experimental findings, SB at sub-MICs strongly suppressed the virulence attributes without significantly attenuating the growth of *V. cholerae*. So, we performed all the experiments at sub-MICs (1/2 MIC and lower) to study the anti-pathogenic role of SB against *V. cholerae.* Additionally, it was also ensured that at a specific optical density (O.D) both SB-treated and SB-untreated *V. cholerae* cultures possess equal number of viable bacteria thereby ruling out the possibility of growth reducing effects of SB on bacteria.

*V. cholerae* is transmitted in humans through the ingestion of the contaminated food and water. Once it enters the host it passes through the acid barrier of the stomach and multiplies in the small intestine. Then in the small intestine it penetrates the mucous barrier and adhere to the microvilli of the epithelial cells (38). After adherence, they start releasing the cholera toxin and this toxin is mainly responsible for the clinical manifestation of this disease (39). Hence, we investigated the anti-CT and anti-TcpA effect in the current study.

Our study revealed that SB impaired the adhering ability of *V. cholerae* to the epithelial cells as well as hindered the expression of cholera toxin (CT). As antibiotic resistance is a serious problem with *V. cholerae*, we checked the effect of SB on two clinical isolates that are resistant to multiple antibiotics. SB was able to inhibit the cholera toxin (CT) production in both the isolates. Our findings are in accordance with earlier studies where various bioactive phytochemicals such as capsaicin, anethole produce anti-virulence effect against *V. cholerae* by inhibiting the cholera toxin production (40, 41). Moreover, the unsaturated fatty acids(UFAs) present in bile and several conjugated forms of UFAs were reported to be potent repressors of cholera toxin in *V. cholerae* (22, 42).

The results obtained from our initial experiments on CT production and adhesion encouraged us to examine the effect of SB at the gene level. We further extended our study to investigate the exact mechanism of action of SB. In the canonical model for *V. cholerae* virulence regulation, the expression of the virulence genes is organized in three-level cascade events. Firstly, AphA and AphB activate TcpPH. TcpPH synergistically acts with ToxRS and activate the transcription of master regulatory gene *toxT* ; and finally ToxT concomitantly activates the transcription of two genes *tcpA* and *ctxAB* that are typically required for colonization and subsequent toxin production (43–45). Quantitative RT-PCR data revealed a significant downregulatory effect of SB on the *ctxAB* and *tcpA* transcripts, however the transcript levels of *toxRS*, *tcpPH*, and the most downstream gene, *toxT*, were all unaffected. The validity of the qRT-PCR data was further confirmed by analyzing the expression of ToxT and TcpA protein levels. Unaltered ToxT level both at RNA and protein level indicates the possibility that inhibition is post-translational. Previous studies on the use of small molecule virstatin, bile against *V. cholerae* were found to exhibit similar effects (21, 22). All these outcomes point that the decrease in CT and TcpA production can be through SB-mediated dysregulation of ToxT binding to their promoter region. This was supported by two crucial Protein-DNA interaction assays (ChIP and EMSA) which demonstrated that SB binds to the ToxT protein and limit its ability to interact with the *tcpA* promoter. Moreover, we tested another butyrate derivative-tributyrate (TB) on ToxT binding. Herein, we observed no interaction between ToxT and TB thus indicating the specificity of SB. Next, the molecular docking was carried out in order to elucidate the interaction between ToxT (target) and SB (ligand). Our docking analysis predicted two important residues Lys 31 and Lys 230 of ToxT that formed conventional H-bonds with the carboxyl group of SB. Earlier works on the crystal structure of ToxT protein when co-purified and crystallized with UFA (cis-palmitoleate) showed that the bound conformation of ToxT is primarily dictated by these lysine-carboxylate interactions (35). Moreover, our docking results displayed a low binding energy (-7.68kcal/mole) between target and ligand indicating that the ligand is in the most favourable conformation. Altogether, these data suggest that SB can interact with ToxT in a manner that can presumably lock ToxT in a “closed” conformation and subsequently prevent DNA binding. These results are in accordance with previous reports which showed decreased ToxT binding to promoter elements by unsaturated fatty acids (22, 42).

We further validated the activity of SB *in vivo* models to show its potential as a future therapeutic in this post-antibiotic era. The effect of SB in *in-vivo* was determined by the suckling mice colonization assay and the rabbit ileal loop model. We investigated the effect of SB on the colonizing ability of *V. cholerae* in both the animal models. The colonizing ability in mammalian host is largely dependent on the expression of TCP, the major subunit of which is TcpA (46–48); in *in vivo tcpA* mutants are considerably less competitive than wild-type strains(49, 50).Our findings demonstrated significant reduction in *tcpA* levels in presence of SB which might explain the possible reason behind decreased colonization of *V. cholerae*. This result corroborates with the previous report where administration of butyrate affected the colonizing ability of *Salmonella* (51, 52). Moreover, SB exhibited a strong inhibitory effect on CT production and subsequently prevented fluid accumulation in rabbit ileal loop. Altogether, these suggest that sodium butyrate was able to abrogate *V. cholerae* colonization in the small intestinal mucosa. There are several reports regarding *in vitro* inhibition of ToxT activity by unsaturated fatty acids and bile but there is still scarcity of validation in *in vivo* models (22, 42).

The hypothesis behind using SB as a therapeutic was that endogenously produced butyrate through fermentative process had a beneficial role in modulating the microbiota by providing energy to intestinal cells, having negative impact on harmful bacteria and regulating the immune related genes of the intestinal system (53–56). Our findings demonstrated similar repressive effect of this compound on the important virulence factors of *V. cholerae*. Moreover, the cytotoxic effect of SB on the human cell line HT-29 was also tested. SB treatment for 24 h at different concentrations (5-80mM) did not show significant toxicity as determined by MTT assay (Figure S2). So, dietary supplementation of this compound during enteric infection by *V. cholerae* or in nutritionally deprived environment could conceivably reduce the bacterial colonization and toxin production without posing detrimental effects on the host system.

In summary, we identified a potential therapeutic candidate namely sodium butyrate (SB), for inhibition of *V. cholerae* virulence gene expression. They function at distinct points in the virulence regulatory cascade by interfering the binding of ToxT to its downstream promoter thereby reducing the activation of CT and TcpA both *in-vitro* as well as *in-vivo* systems and subsequently inhibiting secretory diarrhoea (Fig. 10). A previous study demonstrated the safety of butyrate supplementation for human use (57–59) and parallelly we have also shown that SB possesses negligible cytotoxic effect on HT-29 cell line. The findings of the present study seem to be promising as it identified SB, a novel inhibitor of virulence factors of cholera. Nonetheless, further research is needed to optimize the ability of SB to inhibit diarrheagenic *V. cholerae* and evaluate the long-term efficacy of this potential anti-diarrheagenic compound.

**Fig. 10).**
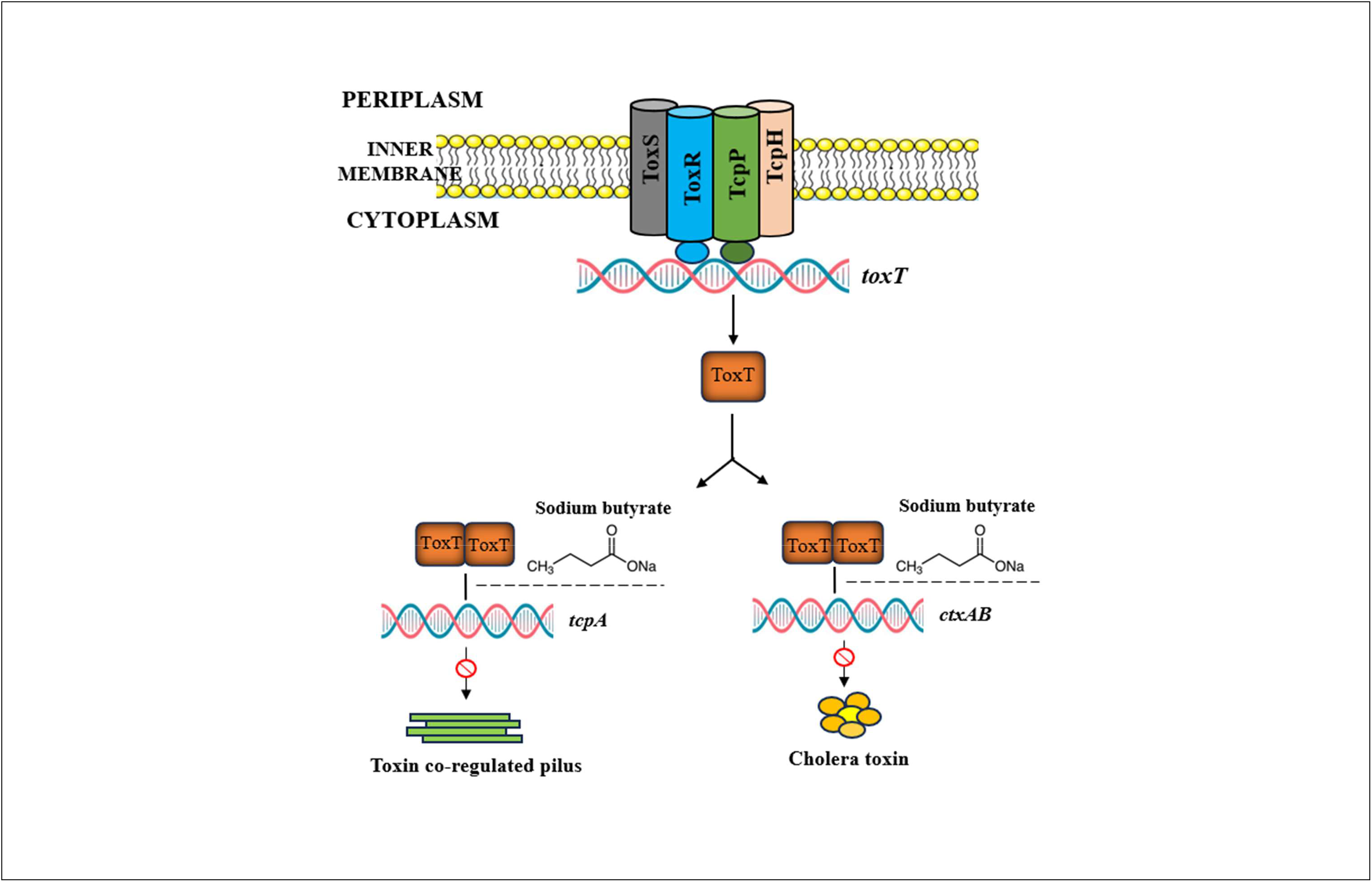
Model showing the inhibition of virulence cascade in *V*. *cholerae* by sodium butyrate. The virulence cascade in *V*. *cholerae* is tightly regulated. TcpPH form an inner membrane complex with ToxR and ToxS to activate transcription of *toxT*. ToxT activates the transcription of *tcpA-F*, which encode the toxin coregulated pilus, and *ctxAB*, which encode the cholera toxin subunits. Based on our experiments, we propose that sodium butyrate inhibits the binding of ToxT protein to the promoter region of *tcpA* /*ctxAB* and eventually inhibits cholera toxin and *tcpA* production.

The findings of sodium butyrate are encouraging as a novel anti-virulent compound and as a potential therapeutic agent for the treatment of *V. cholerae* infection. Although there are several reports regarding butyrate as a promising antibacterial agent, however detailed research on pharmacokinetics relating to human evidence may be worthwhile before SB administration. However, butyric acid has been Generally Recognized as Safe (GRAS) for its use in foods (Butyric acid-21CFR182.60, Food and Drug Administration (Food and Drug Administration [FDA], 2017), but there is a growing body of evidence about the poor bioavailability, unfavourable organoleptic properties such as offensive odour and unpleasant taste (60).There have been multiple reports on butyrate modifications, delivery systems (61, 62) which are showing great efficacy against the targets. Future research will determine the efficacy of these modified products, their delivery system against diarrheagenic bacteria with particular emphasis against *V. cholerae*.

## MATERIALS AND METHODS

### Bacterial strains and culture conditions

Three strains of *V. cholerae* were used. O1 El Tor N16961 was used as a standard strain and two O1 El Tor variant Micro78 [AMP/S/SXT/NA] and BCH13298 [AMP/S/SXT/NA]. All of them were cultured in Luria–Bertani (LB) broth supplemented with 100 μg/ml of streptomycin at 37°C with 180 rpm continuous shaking. Moreover, for CT analysis, bacteria were grown in AKI media containing 0.4% yeast extract (BD Difco, San Diego, CA), 1.5% Bacto peptone (BD Difco, San Diego, CA), 0.5% NaCl, and 0.3% freshly produced NaHCO_3_ (Merck, Burlington, MA) pH 7.2 and at 37°C under static condition.

### Determination of minimum inhibitory concentration (MIC), sub-minimum inhibitory concentration (sub-MIC) and minimum bactericidal concentration (MBC)

The MIC of sodium butyrate against *V. cholerae* N16961, Micro 78, BCH13298 were assessed (63) by microdilution method according to CLSI guidelines (64). The optical density of the bacterial culture was adjusted between 0.08-0.12 value at 600nm that was equivalent to 0.5 McFarland standard turbidity. In a 96-well microtiter plate, 100µL of bacterial suspension (1×10^5^ CFU/mL) was inoculated to each well. Sodium butyrate was dissolved in Muller Hinton Broth (MHB) and combined with the bacterial suspension as required to create a range of concentrations from 5 to 640 mM. The microtitre plate was incubated for 24 h at 37°C. Bacterial growth was assessed by the presence of turbidity and a pellet on the bottom after 24-hour. The concentrations of sodium butyrate that did not affect the bacterial growth was chosen as the sub-MICs for the investigation, while the lowest sodium butyrate concentration that caused no change in OD_600_ after 24 hours was identified as the MIC (65). The lowest concentration of the compound where it can kill 99% of the bacteria was defined as the MBC.

### Vibrio cholerae growth assay

The growth curve of *V. cholerae* El Tor strains were examined using an overnight culture in LB media, followed by the addition of sodium butyrate at various doses 1/8^th^ MIC (10mM), 1/4^th^MIC(20mM), 1/2^th^MIC(40mM), 1xMIC(80mM) and 2xMIC(160mM). Briefly, in 5 mL of LB broth, one colony was inoculated and cultured overnight at 37°C under shaking conditions. The overnight grown stationary phase cells were collected, centrifuged, washed twice in PBS and their number was adjusted to 1 × 10^9^ cells per ml using LB medium. Then in fresh 25 mL LB broth, an initial inoculum of 1 × 10^8^ cells per ml was added and the cultures were maintained at 37°C for a further 24 hours with constant shaking condition (66). The viability of the bacteria was determined by dilution plating onto LA plates treated with streptomycin at different time hours followed by colony count.

### GM1 enzyme-linked immunosorbent assay

GM1 enzyme-linked immunosorbent test (ELISA) was used to measure the quantity of cholera toxin (CT) in the culture supernatant as described by (67–69). Firstly, the cells were grown in AKI media for 18 h at 37°C under static condition. After 18 h, the total bacterial culture was collected and cell-free supernatant (CFS) was prepared by centrifugation of the culture. MaxiSorp ELISA plates (Nunc, Rochester, NY) were used for this assay. Briefly, 96-well polystyrene microtiter plates were coated with GM1 ganglioside overnight. The plates were washed three times with PBS (pH 7.4), 0.2% bovine serum albumin [BSA], and 0.05% Tween 20 and then 1% (w/v) BSA was applied to block the GM1-coated plates for 30 min at room temperature. The plates were then washed three times to remove BSA. Next, CFS was added to the wells and incubated for 1 h at 37°C. Then, 1% BSA was used to block the wells for 30 min at room temperature, and the wells were washed three times thereafter. Subsequently, a goat anti-CT polyclonal antibody (1:1000) and then an AP-linked rabbit anti-goat IgG antibody (1:1000) were added to the wells and allowed to incubate for 1 h at room temperature each; the plates were washed following each step. For development of the CT-antibody complex, we utilized *p*-nitrophenyl phosphate substrate solution (Invitrogen) according to the manufacturer’s protocol. The colour intensity in each well was measured at 450 nm in a plate reader and represented as ng of CT per ml (70). We estimated CT amounts in the samples by comparison to a standard curve This technique was also used to assess the amount of CT secreted in each of the rabbit ileal loop.

### RNA isolation and quantitative RT-PCR

*V. cholerae* cells were grown to mid-logarithmic in the presence or absence of SB to study its effect on the expression of genes related to virulence. Total RNA was extracted from these cells using the Trizol reagent. The extracted RNA was dissolved in RNase-free water and stored at -80°C. To remove any contaminated genomic DNA, the RNA was treated with RNase-free DNase I (Invitrogen). Following the instructions provided by the manufacturer, 1µg of RNA was used from each sample to create cDNA using verso cDNA Synthesis Kit (Thermo Scientific). The mRNA transcript levels were quantified by quantitative PCR (qPCR) by using SYBR green (AB Applied Biosystems) and each gene-specific forward and reverse primers synthesised from IDT (Integrated DNA Technologies). The reaction was performed in 7500 Real-Time PCR detection system (Applied Biosystems). Using *recA* (VC 0543) as an internal control (Table S1), the relative transcription of the target genes were determined using Livak’s 2^-ΔΔC^_T_ technique (71).

### Western blot analysis

Untreated and treated cell lysates were prepared in lysis buffer (10 mM Tris–HCl, pH 8.0, 1 mM EDTA, 1% Triton X-100, 1 mM PMSF, protease inhibitor added) followed by sonication and then centrifugation at 10,000 rpm for 30 min at 4°C. After centrifugation, the supernatant was collected in fresh tubes and the cell pellet was discarded. After protein estimation by Lowry method, lysates were heated in SDS-PAGE sample buffer [SDS (sodium dodecyl sulphate, also called lauryl sulphate), b-mercaptoethanol (BME), bromophenol blue, glycerol, and Tris-glycine at pH 6.8] before being run at 100 volts on a 12.5% SDS-PAGE gel. Then the gels were transferred on to PVDF membrane. The membranes were incubated with primary antibodies overnight at 4°C after being blocked for 1 h at room temperature with 5% skimmed milk diluted in TBST buffer (20 mM Tris-HCl, 150 mM NaCl, 0.1% Tween20). The primary antibodies used are rabbit polyclonal anti-ToxT (Cat#BB-SAP50) (1:5,000), rabbit polyclonal anti-TcpA antibody (Cat#BB-SAP50) (1:8,000). Membranes were then incubated with goat anti-rabbit secondary antibody (dilution 1:10,000) for 2 h at room temperature after primary antibody treatment. Finally, TBST was used to wash the membranes for 30 minutes (72). Bands were developed in the ChemiDoc MP Imaging System (Biorad) using chemiluminescent HRP substrate (millipore)

### Purification of protein

*Escherichia coli strain BL21(DE3*) was used to purify the nickel binding protein (NBP)-ToxT using the pHIS-Tev plasmid harbouring the 6X His-ToxT construct. Briefly the cells were grown at 37°C overnight, then they were sub-cultured in new LB broth in 1:100 ratio and continued to grow there until the optical density at 600 nm (OD_600_) reached 0.5. After that the culture was induced by 0.5 mM of isopropyl-D-thiogalactopyranoside (IPTG) for 3 h at 37°C. ToxT protein was extracted as an insoluble protein from inclusion bodies by sonication and solubilized in 10 ml of buffer A (25mM Tris, 100mM NaCl, 0.1%Triton X-100, 7M Urea pH 7.5).The denatured protein was loaded on top of Ni^2+^-conjugated agarose beads already preequilibrated in buffer A. The column was washed with 20 ml of buffer A and the denatured protein was eluted with 10 mL of buffer A containing 250 mM of imidazole. The fractions containing 6X His-ToxT were collected and then renatured slowly by the gradual removal of urea by dialysis through 10 kD membrane. SDS-PAGE was used to analyze the samples. The amount of protein present in the sample was measured using Lowry’s method.

### Chromatin immunoprecipitation

The effect of SB on ToxT occupancy at *tcpA* promoter was studied by chromatin immunoprecipitation (ChIP) (72). The *V. cholerae* cells were grown in presence of SB (20mM) for 4 h. After incubation, the culture was crosslinked in 1% formaldehyde. Crosslinking was stopped by the addition of glycine to a final concentration of 125 mM, and bacteria were washed thrice in ice-cold PBS buffer. The cells were resuspended in lysis buffer(10mM Tris-HCl pH=8.0, 50mM NaCl containing 20 ng/µl RNase A, 10^5^ kU of ready lyse lysozyme) followed by 30 minutes incubation at 37^0^C. One volume of double strength IP buffer[200mM Tris HCl pH=7.5, 600 mM NaCl, 4% Triton X-100 containing protease inhibitor cocktail and 1 mM phenylmethylsulfonyl fluoride(PMSF)] was added to each lysate and the DNA was sonicated in the range of 200 to 1000 bp. The cell debris was removed by centrifugation for 10 min at 12,000 rpm,4°Cand a fraction of the supernatant was stored as the input sample for IP assays. The nucleoprotein complexes were immunoprecipitated by overnight incubation at 4°C with 8 μg anti-ToxT antibody. Salmon sperm DNA treated G agarose beads (Thermo Scientific), pre-washed with lysis buffer was used to pull down the antibody protein DNA complexes for 4 h at 4°C. Beads were then washed once with lysis buffer, once with wash buffer (500 mM NaCl lysis buffer), once with LiCl immune complex buffer [20mMTris–HCl pH8.0, 0.25MLiCl, 1mM EDTA,0.5% (wt*/*vol) NP-40, 0.5% (wt*/*vol) DOC, 1% (wt*/*vol) PMSF], and once with buffer TE (10 mM Tris–HCl pH 8.0, 1 mM EDTA). The immunoprecipitated complexes (IP) were eluted from the beads by incubation at 65°C for 30 minutes in TE buffer containing 1%SDS. Reversal of cross-linking was done by incubation with 20µg of proteinase K for 1 hour at 65◦C. Both control input and immunoprecipitated DNA samples were purified using a PCR purification kit(Qiagen) and eluted in 30 µl DNase free water. Real-time quantitative PCR (qPCR) was used to quantitate promoter occupancy by ToxT as formerly described (73) using the primers EMSA TcpA FP and EMSA TcpA RP (Table S1). The amount of immunoprecipitated DNA (IP DNA) was estimated as a percentage of the input DNA by using the formula IP = 2(^CT.^_input_ ^-CT^_IP_), where CT is the fractional threshold cycle of the input and IP DNAs. The promoter DNA was also analyzed by agarose PCR gel electrophoresis using the primers EMSA TcpA FP and EMSA TcpA RP(Table S1).

### Electrophoretic mobility shift assay

The chromosomal DNA of *Vibrio cholerae* N16961 was used as a template in the PCR process to amplify DNA fragment of size 150 bp for *tcpA* promoter region (73). Then this amplified product was labeled with biotin at 5’ end followed by purification using the Qiaquick nucleotide removal kit (QIAGEN).To check the drug-protein interaction, the binding reactions were set up where purified His-tag ToxT protein was pre-incubated with SB for 20 minutes in binding buffer (10 mM Tris-HCl [pH 7.5], 100 mM KCl, 1 mM EDTA, 1 mM dithiothreitol, 200 μg of bovine serum albumin per ml, 10% glycerol) followed by the addition of fixed amount of biotinylated-labelled DNA fragment(1µM) and incubated for another 20 minutes. Another butyrate derivative tributyrate (TB) was used to check the specificity of the interaction.

Further, to check possible drug-DNA interaction, labelled DNA (0.5µM) was pre-incubated with SB for 20 mins in binding buffer (74). After incubating for 20 mins, purified ToxT protein (18 µM) was added, and the incubation continued for another 20 mins. The samples were loaded on 4% native polyacrylamide gel and electrophoresed at 100 V, 4°C in 0.5X Tris-borate-EDTA buffer. DNA was then transferred from gel to charged nylon for nucleic acid blotting (Millipore) followed by cross-linking using a UV Stratalinker from Stratagene and detected using LightShift™ Chemiluminescent EMSA Kit (Thermo Scientific™

### Binding curve analysis.

Briefly, to determine the K_D_ (equilibrium dissociation constant) for samples containing sodium butyrate (SB) and compared to protein bound to DNA without inhibitor, the percentage of protein bound with labeled DNA was determined for each lane. This was then fit to the following equation: percent 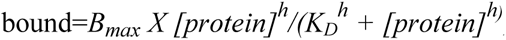, where *h* is the Hill coefficient and *B_max_* is the amount of bound DNA at which the curve plateaus, which was set to a constraint of 100% using GraphPad Prism 9 software. The *K_D_* values for each condition were compared to each other using the extra sum of squares F test to determine if the two values were statistically different.

### Molecular docking

The 3D structure of the docking target ToxT was downloaded from the RCSB Protein Databank Server and visualized using UCSF Chimera version 1.11. The said protein was prepared for docking using Yasara engine and then saved as a new PDB file using Chimera. The 3D structure of the ligand Sodium butyrate was obtained from Pubchem (Compound CID: 5222465) in SDF format and converted to PDB file using UCSF Chimera (75)

The Patchdock-Firedock online docking servers were used for molecular docking. The docked complex was visualized using PyMOL version 1.7.2.1 and the results represented (76)

### Cell culture

To study the anti-adhesion effect of SB in cell culture conditions, HT-29 cells were used. The cells were maintained in Dulbecco’s modified Eagle’s medium (DMEM) (Sigma-Aldrich, St. Louis, MO). The media was additionally supplemented with 10% heat-inactivated fetal bovine serum (Sigma, USA) along with 1% non-essential amino acids and penicillin/streptomycin (Himedia, India). The cells were kept and maintained at 37°C in a humidified 5% CO_2_ incubator (77)

### Adhesion assay

Adhesion of *V. cholerae* to HT-29 cells was done according to previously described method (66, 70)In 12-well cell culture plates, HT-29 cells were splitted and grown till 80% confluency. The *V. cholerae* cells were grown upto mid-log phase and 1.2 ×10^8^ bacteria (1: 100 MOI) were inoculated to HT-29. SB at varying concentrations (20, 40 and 80 mM) was mixed with DMEM in each well. The adhesion assay was performed by the incubation of the bacteria /cell for 1 h at 37°C. To remove the unadhered bacteria the cells were washed thrice with PBS pH=7.4 followed by the detachment of the adhered bacteria using 0.1% Triton X-100.The cells were then serially diluted and counted by plating them onto LB agar plates supplemented with 100 μg /ml streptomycin*. In vivo* adhesion of *V. cholerae* along the rabbit intestinal cells was also evaluated. After 18 h of the experiment, intestinal loop sections were recovered, washed in PBS followed by homogenization and serial dilution. The count of adherent bacteria was enumerated by plating them on LB agar plates supplemented with streptomycin (100 μg /ml).The percentage of adherent bacteria (%adherent bacteria) was calculated using the following equation, Adherent bacteria 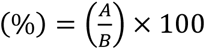; where A = No. of adhered bacteria, B = Total no. of recovered viable bacteria.

### MTT Assay

In 96 well culture plates, HT29 cells were grown at a density of 1×10^4^ cells per well. After reaching confluency, cells were treated with different doses of SB (5-80mM). Cells were incubated at 37°C for 24 h in a 5% CO_2_ incubator. Cellular viability was determined by using Colorimetric Cell Viability Kit IV (MTT) (Promokine). 20 μl of the reagent was added and incubated at 37°C for 4 h. The purple crystal formazan was then solubilized by the addition of 300 μl of DMSO. At 570 nm, OD was measured and % viability was calculated and represented graphically.

### Suckling mice colonization assay

The intestinal colonization assay by *V. cholerae* was performed in suckling mice according to the previously described protocol (78). Briefly, 4 to 5-day old suckling BALB/c mice were removed from their mothers 1 hour just before inoculation. Each mouse was then orogastrically fed with 1×10^5^ CFU bacterial culture with or without compound (SB) and the mouse fed only with PBS served as negative control. Each mouse was maintained at 30°C without their mother. They were then sacrificed after 18 h of infection, whole intestines were collected, washed with PBS and homogenised mechanically. Serial dilution of intestinal homogenates was done and plated onto LB agar petri plates supplemented with 100 μg/ml streptomycin to calculate the recovered CFUs. From each of the samples the bacterial count was measured to ascertain their impact on bacterial colonization in intestinal epithelial cells.

### Ligated rabbit ileal loop assay for fluid accumulation and bacterial recovery

Rabbit ileal loop experiments were performed in New Zealand white rabbit (1.5 to 2 kg) as previously described (79). Prior to surgery, the rabbits were starved for 48 h and just given water as needed. The animals were anesthetized by intramuscular injection of xylazine (5 mg/kg of body weight) and ketamine (35 mg/kg of body weight). A surgical incision was made in the wall of the abdomen, and then the ileum was cleaned and ligated into separate loops of about 10 cm. Each loop was injected with 1×10^9^ CFU per ml *V. cholerae* N16961 with or without SB (at 20, 40 and 80mM) and loop injected only with PBS served as negative control loop. After 18 h, the animals were euthanized, and the loops were removed. The length of each loop and the volume of the accumulated fluid were recorded and accumulated fluid per unit area (F/A) was represented as loop fluid volume (ml)/length (cm) ratio (66, 70). The collected fluid was also used to estimate the CT production in the intestine.

### Histopathological study

Rabbit ileal tissue samples were fixed with 10% buffered formalin. Gradient dehydration with ethanol (50–100%) was performed, followed by xylene treatment. Tissues were then embedded in paraffin (56–58°C) at 58 ± 1°C for 4 h. After deparaffinization with xylene, the staining was done with haematoxylin-eosin (H and E) and mounted in DPX under a clean coverslip. Histological variations were visualized using a bright field microscope (Motic, Germany), and images were captured at different magnifications.

### Animal ethics

Animal experiments were carried on following the guideline proposed by the Committee for the Purpose of Control and Supervision of Experiments on Animal (CPCSEA), Government of India. All animal experiments performed here were subjected to the approval (registration no: 68/GO/ReBi/S/99/CPCSEA) by the Institutional Animal Ethics Committee of the National Institute of Cholera and Enteric Diseases.

### Statistical analysis

The experiments were performed in triplicates. All data were represented as mean ± standard error of mean (S.E.M) and analyzed using GraphPad Prism 9.0 software. The difference in statistical level was analyzed using unpaired (Student’s) t test, one-way ANOVA (Tukey’ s multiple comparison tests) and two-way ANOVA. A *P* values *<* 0.05 implies statistical significance. Significant levels are denoted as ∗for *P <* 0.05, ∗∗for *P <* 0.01, ∗∗∗for *P <* 0.001 and ∗∗∗∗for *P <* 0.0001.

## ACKNOWLEDGEMENTS

SK thanks the University Grants Commission (UGC). PM thanks the Council for Scientific and Industrial Research (CSIR) and PH thanks Indian Council of Medical Research (ICMR) for supporting with fellowship. We also thank Indian Council of Medical Research (ICMR) for institutional funding.

